# The *E. coli* NudL enzyme is a Nudix hydrolase that cleaves CoA and its derivatives

**DOI:** 10.1101/2020.01.31.929182

**Authors:** Joseph Rankin Spangler, Faqing Huang

**Author notes:** E-mail addresses (Joseph Rankin Spangler), (Faqing Huang).

## Abstract

The process of bacterial coenzyme A (CoA) degradation has remained unknown despite the otherwise detailed characterization of the CoA synthesis pathway over 30 years ago. Numerous enzymes capable of CoA degradation have been identified in other domains of life that belong to the Nudix superfamily of hydrolases, but the molecule responsible for this process in the model bacterial system of *E*. coli remains a mystery. We report here that *E*. coli contains two such Nudix enzymes capable of CoA degradation into 4’-phosphopantetheine and 3’,5’-adenosine monophosphate. The *E*. coli enzymes NudC and NudL were cloned in various promoter-fusion constructs in order to purify them as soluble active enzymes and characterize their ability to catalyze the phosphohydrolysis of CoA. NudC, an enzyme known to hydrolyze NADH as its principal substrate, demonstrated the ability to hydrolyze CoA, among other coenzymes, at comparable rates to eukaryotic Nudix hydrolases. NudL, a previously uncharacterized enzyme, demonstrated the ability to cleave only CoA and CoA-related molecules at a rate orders of magnitude slower than its eukaryotic orthologs. NudC and NudL therefore represent a previously uncharacterized pathway of CoA degradation in the highly studied *E*. coli system. While the two enzymes display some substrate overlap, their respective activities imply that NudC may play a role as a general coenzyme hydrolase, while NudL specifically targets CoA. These data further suggest a role for these enzymes in the regulation of bacterial CoA-RNA.

## 1. Introduction

The Nudix enzymes are a superfamily of phosphohydrolases that exist in all domains of life [1] and cleave nucleoside diphosphates linked to another moiety [2, 3]. Originally characterized to function in cellular housecleaning for the removal of deleterious oxidized nucleotide derivatives [4], a number of coenzyme-specific Nudix enzymes have been characterized over the years that extend the role of the superfamily to include coenzyme turnover [5–8]. Structural studies of Nudix enzymes have resulted in an amino acid sequence motif known as the Nudix box (Gx_5_Ex_5_[UA]xREx_2_EExGU, Prosite 00893) [4] that has since been used to identify purported members of the Nudix superfamily across different phyla. Further studies into coenzyme-targeting Nudix enzymes have uncovered coenzyme-specific motifs such as LLTxR[SA]x_3_Rx_3_Gx_3_FPGG (Prosite UPF0035) that gives coenzyme A (CoA) specificity [8] and SQPWPxPxS for reduced nicotinamide adenine dinucleotide (NADH) specificity [6] that have been used to predict substrate preference [2]. Such sequence motif criteria has led to various studies characterizing CoA hydrolase orthologs in yeast [8], bacteria [9], nematodes [7], plants [10] and mammals [11] capable of cleaving the phosphoanhydride bond of CoA to produce 4’-phosphopantetheine (pPan) and 3’,5’-adenosine diphosphate (3’,5’-ADP).

The bacterial CoA synthesis pathway has been almost completely characterized since the 1980s when it was a hot area for the development of antibacterial drugs [12–14]. Using *E. coli* as a model organism, both the forward and reverse directions of the pathway were demonstrated via reversible activities of the respective enzymes catalyzing each step [15] with the peculiar exception of CoA degradation itself. While investigators have discovered and characterized Nudix superfamily enzymes with such capability in a diverse set of organisms, the *E. coli* counterpart has remained elusive only to be hypothesized as the gene product named YeaB or NudL [2]. Such a lack of information is interesting considering the monumental discoveries of coenzyme-linked RNA in that same organism [16, 17], which brings into question both synthesis and degradation of such molecules. Indeed, the degradation of NAD- [18] and CoA-linked RNA [19] in *E. coli* has been hypothesized to be carried out by the Nudix hydrolase NudC, but sequence and structural evidence insinuates the activity of NudC is that of a general coenzyme hydrolase [5, 20]. An enzyme with specific activity for CoA hydrolysis in *E. coli* therefore remains hypothesized but unknown.

Here we demonstrate for the first time that the *E. coli* protein NudL exhibits phosphohydrolase activity towards CoA and its derivatives as predicted by its sequence similarity to other known CoA-hydrolyzing orthologs. We also show evidence that NudL is a CoA-specific enzyme incapable of displaying observable activity towards structurally diverse substrates. Considering that NudC has never been characterized with CoA as a substrate, comparisons of NudL and NudC establish a hierarchy of activity between the two enzymes.

## 2. Materials and Methods

### 2.1 Materials

All chemicals were purchased from Sigma-Aldrich (St. Louis, MO) unless otherwise stated. DNA oligos were purchased from IDT (Coralville, IA). Competent cells were obtained from New England Biolabs (Ipswich, MA) and Lucigen (Middleton, WI). Enzymes were stored in 40% glycerol at −20 °C until use.

### 2.2 Cloning Constructs

Cloning constructs were assembled by *in vivo* homologous recombination in DH5α and High-Control 10G cells following the protocol described elsewhere (Huang & Spangler, submitted for review). Briefly, NudC and NudL genes were amplified from the *E. coli* genome by polymerase chain reaction (PCR) with Hot Start Q5 (NEB) and primers flanking the open reading frames that added 18 base pairs of homology to pMBP-Parallel, pETite-nHis-SUMO, or pET-28a backbones that were separately linearized by PCR. Nudt7 was prepared similarly from a cDNA library from murine C3H cells as a gift from Yan-Lin Guo. In the case of pMBP-Parallel, PCR amplification of the backbone was carried out to both linearize the vector for recombination in addition to replace the recognition site of TEV protease with thrombin for the downstream removal of the maltose-binding protein fusion. Backbone and potential insert were combined in estimated 1:1 volume ratio in 10 μL aliquots of competent cells and transformed via heat shock according to manufacturer’s protocol. Constructs were checked by Colony PCR for correct size using general backbone primers and confirmed to be inserted in-frame by Sanger sequencing (Eton Bioscience, Research Triangle, NC).

### 2.3 Protein Expression and Purification

Proteins encoded on pETite-nHis-SUMO backbone were expressed in High-Control Bl21(DE3) cells (Lucigen), while those encoded on pET-28a and pMBP-Parallel were expressed in Bl21(DE3) cells (NEB). Cultures containing constructs of NudC, NudL, or Nudt7 fused with N-terminal SUMO (pSUMO-NudL, pSUMO-NudC, pSUMO-Nudt7), maltose-binding protein (pMBP-NudL), or His_6_-tag alone (pET28a-NudL) were grown at 37 °C in Luria Bertani broth (10 g/L tryptone, 5 g/L yeast extract, 10 g/L NaCl) to an OD_600_ of 0.5 and induced with 0.2 mM IPTG at 16 °C shaking overnight. Cells were harvested at 10,000 *x g* with a JLA 25.50 rotor (Beckman) in an Avanti J-26 XP centrifuge (Beckman Coulter), cell pellets were frozen at −20 °C for at least 1 hour and resuspended in phosphate buffer (250 mM phosphate pH 8, 0.5 M NaCl) before sonicating on ice using probe tip XL-2000 sonicator (Misonix) for 10 second bursts with 1 minute rest. Cell lysate was clarified at 40,000 *x g* at 4 °C with a JLA 16.250 rotor and supernatant was purified by affinity chromatography with Ni-NTA resin (Thermo) for His_6_-tagged proteins or Amylose resin (NEB) for MBP-tagged proteins.

Proteins containing His_6_-tags were purified by binding to Ni-NTA resin and washing with phosphate buffer containing 20 mM imidazole while monitoring protein absorbance at 280 nm. Protein was eluted with 300 mM imidazole and fractions with the highest absorbance were pooled for dialysis against a storage buffer (20 mM phosphate pH 8, 150 mM NaCl) to remove the imidazole. A similar protocol was carried out for MBP-tagged proteins using amylose resin, where target protein was eluted with 50 mM maltose. Eluate fractions were combined using an M50 centrifugal filtration column (Amicon) to reduce volume and salt to 20 mM phosphate (pH 8) and 150 mM NaCl. Desalted enzyme fractions and dialyzed enzymes were supplemented with 40% glycerol for storage at −20 °C and quantified by absorbance at 280 nm using extinction coefficients calculated by ExPASy (www.expasy.org).

### 2.4 CoA-RNA Preparation

CoA-RNA was prepared following a previously described method [21]. Briefly, *in vitro* transcription using the Epicentre Ampliscribe T7-Flash kit (Madison, WI) under control of the φ2.5 promoter was carried out in the presence of dephospho-CoA (dep-CoA) and α-^32^P-ATP following manufacturer’s protocol to generate internally radiolabeled CoA-RNA with a length of 10 nucleotides. CoA-RNA was purified from 5’-triphosphate RNA using thiopropyl Sepharose 6B resin following a procedure described elsewhere [21, 22] and visualized for purity by 12% denaturing PAGE. CoA and CoA-RNA dimers were prepared by natural oxidation at room temperature over time as determined by high pressure liquid chromatography (HPLC) or polyacrylamide gel electrophoresis (PAGE), respectively.

### 2.5 Enzyme Analysis

Enzyme activity assays were carried out by varying concentrations of micromolar enzyme and millimolar substrate in 50 mM Tris (pH 7.5) with 10 mM MgCl_2_ at 37 °C. Reaction aliquots were analyzed monitoring absorbance at 260 nm and retention time compared to standards via HPLC using an Alltech Allsphere SAX 5u 250×4.6mm column (Deerfield, IL) at 1.5 ml/min with 40 mM KH_2_PO_4_. Chromatograms were integrated and converted to reaction velocities to fit velocities vs. substrate concentrations using Origin software (Northampton, MA). Reactions of enzyme vs. CoA-RNA were carried out similarly, but incubated at 37 °C for 20 minutes before analyzing reaction progress by 12% denaturing PAGE containing 7 M urea. After electrophoresing 10 minutes, gels were dried and exposed for visualization by phosphorimaging (Molecular Imager; Bio-Rad Laboratories, Hercules, CA).

## 3. Results

The 579 bp gene encoding NudL was cloned into the pET-28a vector for recombinant expression under the control of the T7 promoter, but the solubility of the 192 amino acid enzyme was found to be poor (Fig. 1A). The typical recombinant expression tricks of lowering induction temperature to 16 °C, reducing inducer concentration to submillimolar, and inducing expression in late log phase for a shorter time could not remedy the insolubility of the construct. As the solubility of recombinant proteins is often aided by fusion to a soluble expression partner such as the yeast chaperone SUMO [23], the DNA encoding NudL was prepared for cloning directly downstream of the SUMO fusion partner in the pETite-nHis-SUMO vector. The resulting construct remained insoluble despite the presence of SUMO (Fig. 1B). The T7 promoter can be desirable in recombinant expression for high protein yield [24], but this high activity seemed to contribute to the insolubility of NudL. Therefore NudL was cloned into the pMBP-Parallel vector (pMBP-NudL) downstream of the highly soluble Maltose-Binding Protein fusion partner where expression would be controlled by the less active *tac* promoter. The 64.5 kD MBP-NudL expression construct was observed to be soluble after sonication (Fig. 1C) with a final yield of 4 mg/L (Figs. 1D and S1). The cloning and expression of NudC was by comparison a simpler undertaking. The 774 bp gene encoding NudC was similarly cloned into the pETite-nHis-SUMO vector (pETite-SUMO-NudC), which was used to express the soluble 41.9 kD SUMO-NudC fusion in a final yield of 29 mg/L (Figs. 1E, S2, and S3).

**Figure 1.**
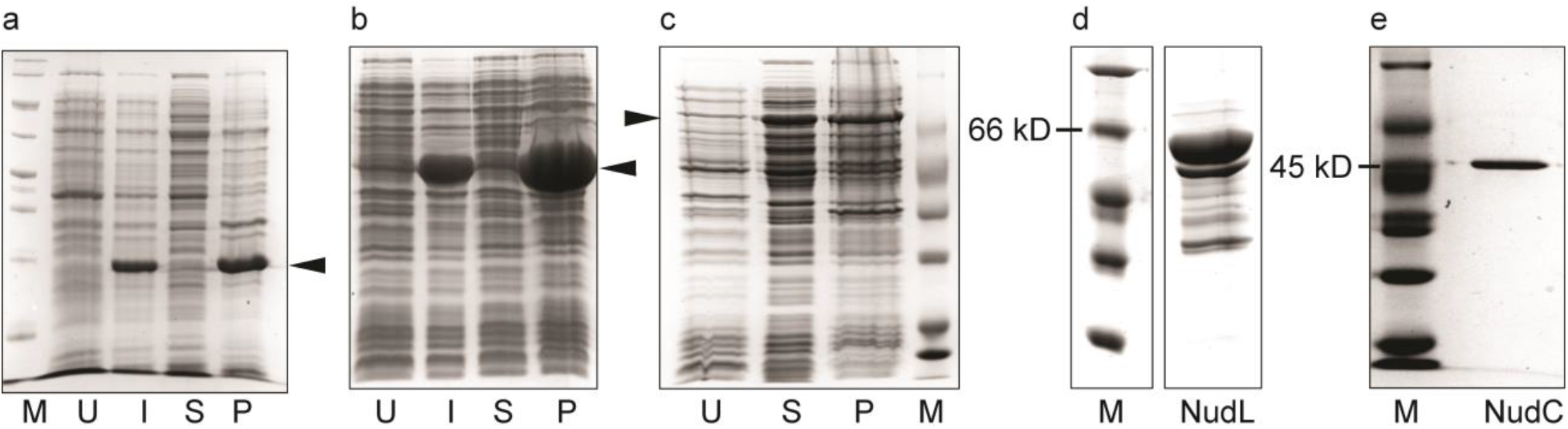
Solubility and purification of NudL and NudC. SDS-PAGE monitoring solubility and purity of recombinant proteins. **A)** Solubility of NudL comparing uninduced cell lysate (U), induced cell lysate (I), soluble fraction of lysate (S), and pellet fraction of lysate (P) with protein molecular weight ladder (M). **B)** Solubility of SUMO-NudL comparing uninduced lysate (U), induced lysate (I), soluble fraction of lysate (S), and pellet fraction of lysate (P). **C)** Solubility of MBP-NudL comparing uninduced lysate (U), soluble fraction of lysate (S) and pellet fraction of lysate (P) with protein molecular weight ladder (M). **D)** Purified MBP-NudL. **E)** Purified SUMO-NudC.

The hydrolysis of CoA (Fig. 2A) can be observed with HPLC by monitoring the 260 nm absorbance of the adenosine base present in both 3’,5’-ADP and CoA. Upon hydrolysis of CoA to form 3’,5’-ADP and 4’-phosphopantetheine, the two molecules should appear as separate peaks eluting from SAX with approximately 3 minutes difference in retention time due to the number of phosphates on each (Fig. 2B). When incubating 4 μM MBP-NudL with 1 mM CoA at the specified reaction conditions, the appearance of two distinct peaks with retention times matching those of CoA and 3’,5’-ADP standards indicated that NudL was capable of phosphohydrolysis. The same activity was observed when incubating MBP-NudL with oxidized CoA, dep-CoA, and oxidized dep-CoA (data not shown) indicating NudL can additionally hydrolyze CoA-derivatives. The hydrolysis of CoA is further dependent on the presence of a divalent metal cofactor, namely Mg^2+^, Zn^2+^, or Mn^2+^, for the removal of the cofactor from reaction mixtures yielded no activity (data not shown).

**Figure 2.**
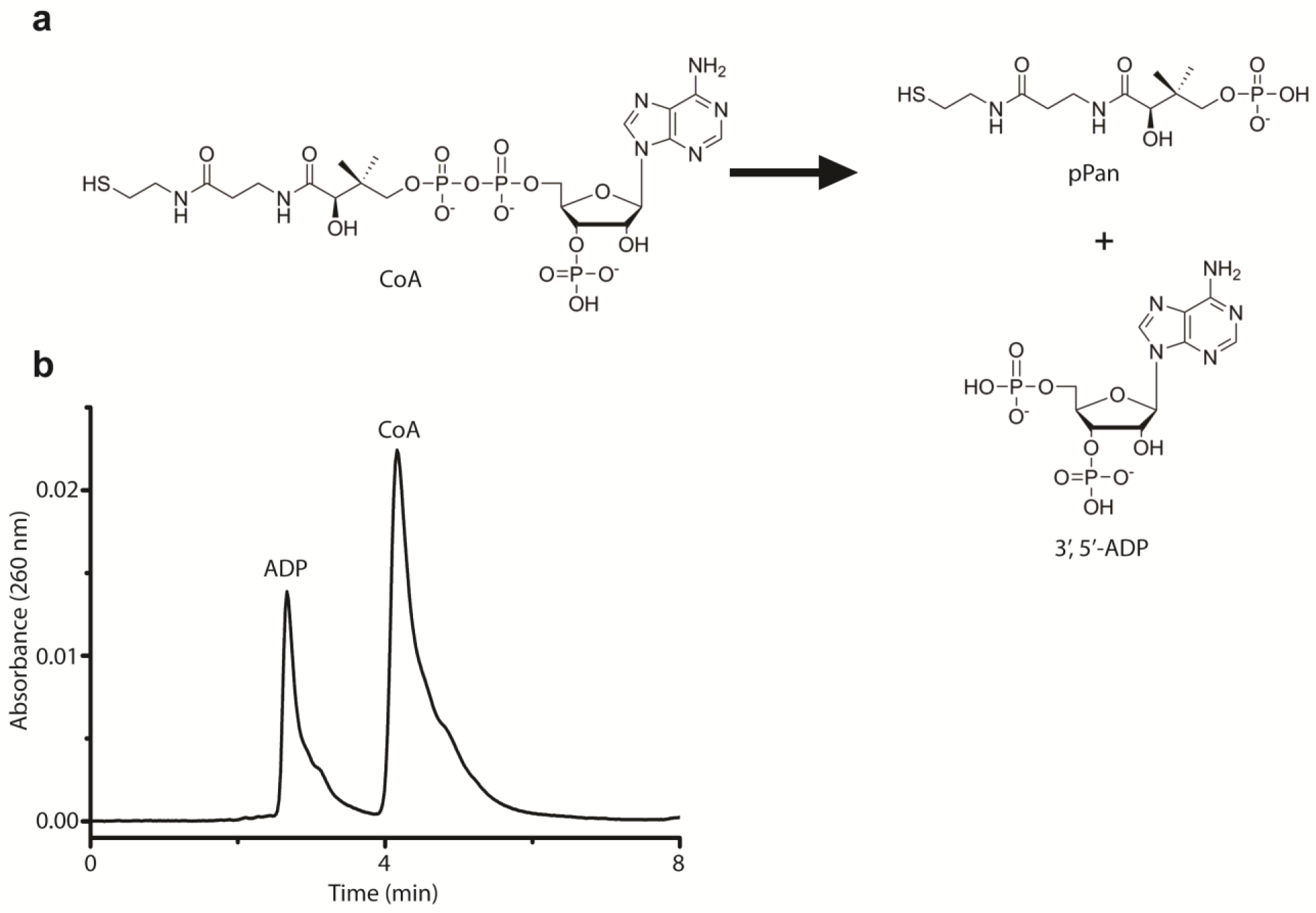
Analysis of CoA hydrolysis. **A)** Chemistry of Nudix-catalyzed hydrolysis of coenzyme A (CoA) forming 4’-phosphopantetheine (pPan) and 3’,5’-adenosine diphosphate (3’5’-ADP). **B)** HPLC chromatogram of Absorbance at 260 nm vs. time showing separation of CoA from 3’5’-ADP.

The quick clearance from the column and distinct peak formation of the products and substrates via HPLC made it convenient to study the kinetics of CoA hydrolysis by injecting reaction aliquots over time (Figs. 3A and 3C). The MBP-NudL fusion was mixed at 1 μM with CoA concentrations between 0.1 and 2.5 mM and incubated with 10 mM MgCl_2_ at 37 °C with injections occurring approximately every 10 minutes (Fig. 3A). The observed reaction velocities were plotted against substrate concentration to generate a Michaelis-Menten plot (Fig. 3B) and calculate the turnover number (k_cat_) of 0.01 s^-1^ and a catalytic efficiency (k_cat_/K_M_) of 0.02 mM^-1^•s^-1^ (Table 1). Considering the slow turnover number obtained, we hypothesized that the MBP fusion might be impeding the activity of NudL. Therefore the MBP fusion was removed by thrombin digestion (Fig. S4). Upon digestion, however, the hydrolysis of CoA could not be observed under the previous reaction conditions (data not shown). Although the removal of solubility-enhancing fusion partners has been known to affect the solubility of a target protein [25], the inability for NudL alone to present observable cleavage of CoA is troubling. Nevertheless, hydrolysis of CoA by the soluble MBP-NudL fusion demonstrates the capability of this enzyme to act on CoA and CoA-derivatives.

**Figure 3.**
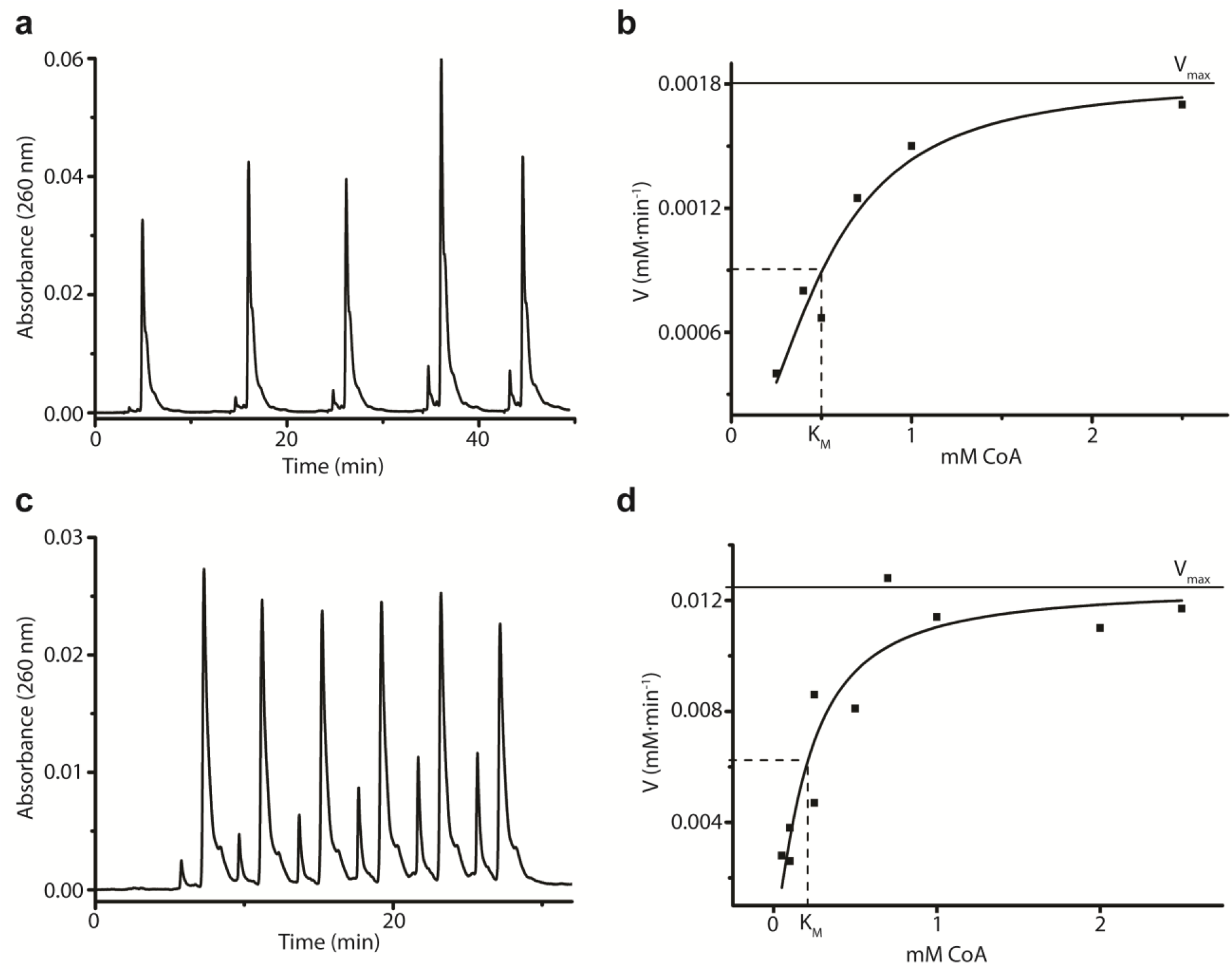
Kinetic Assays of MBP-NudL and SUMO-NudC with CoA. **A)** HPLC chromatogram obtained when combining MBP-NudL and CoA at 37 °C and injecting aliquots every 10 minutes. **B)** Integration of HPLC chromatograms of MBP-NudL vs. CoA were used to produce velocities, which were then plotted against substrate concentration to calculate K_M_ and V_max_ and subsequently used to calculated k_cat_. **C)** HPLC chromatogram obtained when combining SUMO-NudC and CoA at 37 °C and injecting aliquots every 5 minutes. **D)** HPLC chromatograms of SUMO-NudC vs. CoA were integrated to produce velocities, which were then plotted against substrate concentrations to calculate K_M_ and V_max_ and subsequently used to calculated k_cat_.

**Table 1.** Kinetic constants of Nudix enzymes acting on CoA.

| Enzyme | $K_M$ (mM) | $k_{cat}$ ($s^{-1}$) | $k_{cat}/K_M$ ($mM^{-1} \cdot s^{-1}$) |
| --- | --- | --- | --- |
| MBP-NudL | 0.51 (*0.09*) | 0.008 | 0.016 |
| SUMO-NudC | 0.21 (*0.07*) | 0.21 | 0.98 |
| Nudt7* | 0.24 | 3.8 | 16 |
| NDX8** | 0.22 | 13.8 | 64 |

Note: Calculated values for MBP-NudL and SUMO-NudC shown with standard error measurements italicized in parentheses compared to values of previously characterized eukaryotic orthologs from mice (\*Gasmi & McLennan, 2001) and worms (\*\*AbdelRaheim & McLennan, 2002).

The activity of SUMO-NudC was tested by combining 4 μM SUMO-NudC with 1 mM NAD^+^ in the presence of 10 mM MgCl_2_ resulting in the elution of separate peaks by HPLC corresponding to adenosine monophosphate and nicotinamide mononucleotide (Fig. S5). Further experiments with Flavin adenine dinucleotide (FAD) showed similar results (Fig. S6), verifying that the enzyme was capable of hydrolyzing structurally distinct coenzymes as previously described [5]. NudL, by comparison, showed no activity towards NAD^+^ or FAD (data not shown). When NudC was combined with CoA, an HPLC elution profile identical to NudL indicated the enzyme follows the characteristic Nudix mechanism for CoA hydrolysis (Fig. 2A). Monitoring reactions of 1 μM SUMO-NudC with 0.1 to 2.5 mM CoA over time resulted in a calculated k_cat_ of 0.21 s^-1^ and a catalytic efficiency of 0.98 mM^-1^•s^-1^ (Figs. 3C, 3D, and Table 1). Taken together, these results reinforce previous observations that NudC acts on a variety of structurally distinct adenosine-containing molecules [5], and is capable of hydrolyzing CoA at a comparable enzyme efficiency to previously characterized Nudix CoA hydrolases [11].

NudC was recently shown to cleave *in vitro* both NAD- [18] and CoA-conjugated RNA [19] endogenous to *E. coli* [16, 17]. Considering that NudL has been shown here to specifically cleave only CoA and CoA-related structures, but not its structurally distinct coenzyme relatives of FAD and NAD, we were curious to see whether NudL would be capable of acting on CoA-RNA. Internally labeled 10 nt CoA-RNA was combined separately with 4 μM MBP-NudL, 4 μM SUMO-NudC, or 10 μM SUMO-Nudt7 at 37 °C for 20 minutes which was included due to its characterized activity against bulky CoA derivatives such as thioesters [11]. The purified CoA-RNA was converted to the oxidized dimer (CoA-RNA)_2_ with time and as a result showed a significant gel shift compared to the monomer CoA-RNA and the phosphohydrolyzed product pRNA (Fig. 4). As predicted, Nudt7 was capable of cleaving both oxidized and reduced CoA-RNA to produce a single band of 10 nt single-stranded pRNA (Figs. 4B and 4C). Despite its oxidation, dimerized CoA-RNA was observed to be cleaved by NudC (Fig. 4A) in a similar pattern to that of Nudt7. NudL, on the other hand, did not display such a pattern (Fig. 4A and 4B). Considering the possibility that NudL could react with reduced CoA-RNA but not oxidized (CoA-RNA)_2_, the reaction was repeated in the presence of DTT to generate the CoA-RNA monomer as a substrate, but no such CoA-RNA cleavage could be observed (Figs. 4C).

**Figure 4.**
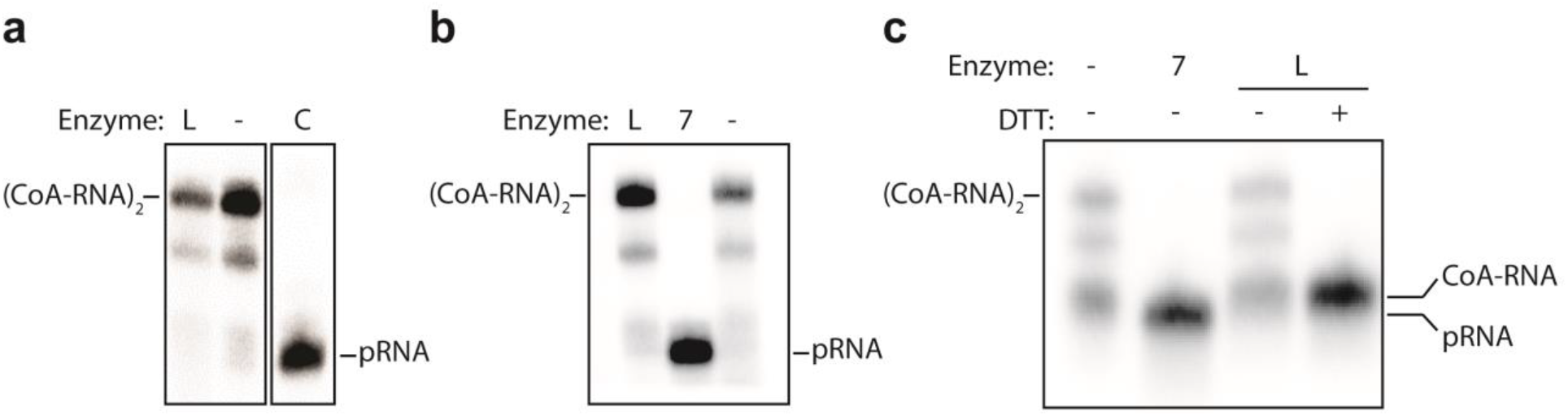
CoA-RNA hydrolysis by Nudix enzymes. Internally labeled 10 nt CoA-RNA was combined in reaction buffer with Nudix enzymes at 20 minutes at 37 °C and then separated by 12% PAGE with single nucleotide resolution to distinguish between CoA-RNA, the naturally oxidized dimer (CoA-RNA)_2_, and the pRNA product of phosphohydrolysis. **A)** Reaction of CoA-RNA with 4 μM MBP-NudL (L), 4 μM SUMO-NudC (C), or no enzyme (-). **B)** Reaction of CoA-RNA with 4 μM MBP-NudL (L), 10 μM SUMO-Nudt7 (7), or no enzyme (-). **C)** Reaction of CoA-RNA in the presence of 50 mM DTT using 4 μM MBP-NudL (NudL), 10 μM SUMO-Nudt7 (Nudt7), or no enzyme (-).

## 4. Discussion

Here we demonstrate that NudL is the CoA specific Nudix hydrolase in *E. coli*. The recombinant expression of NudL proved to be a difficult undertaking due to its solubility *in vitro*. The inability for SUMO to influence the solubility of the enzyme was perplexing, as it is a popular solution to expression problems with other investigators [25] as well as with our own constructs. The MBP did succeed in solubilizing NudL, but its incorporation was problematic due to its large size of over 40 kD which is nearly twice that of NudL. The size of MBP is partly what makes it a great solubility-enhancer, but it seemed to have impeded the observable enzyme activity of NudL. While NudL activity was demonstrated with the attached fusion protein, the calculated k_cat_ and catalytic efficiency were drastically lower than its eukaryotic counterparts (Table 1). With a turnover number over 400- and 1700-fold lower than murine Nudt7 [11] and *C*. elegans NDX8 [7], respectively, it seemed that the MBP fusion may have altered the kinetics of the NudL enzyme. The removal of the fusion, however, resulted in a loss of observable activity, either due to solubility changes in the absence of the fusion or extended incubation at 37 °C during protease digestion. Protease treatment at room temperature, however, resulted in a similar lack of NudL activity, implying that the enzyme had lost stability without its fusion partner. The solubility issues of NudL bring into question the enzyme’s cellular function, and whether or not it exhibits *in vivo* activity at a detectable level.

The kinetic studies with CoA proved SUMO-NudC to be a much faster enzyme than MBP-NudL. SUMO-NudC displayed a turnover number more than 20-fold higher than MBP-NudL, and only an order of magnitude slower and less efficient than the murine Nudt7 (Table 1). The enzyme has been previously characterized to hydrolyze a wide variety of nucleoside-derived substrates [5, 18, 19, 26] with optimal activity for NADH, and we observed an 18-fold lower turnover number for CoA than for NADH [5]. These results imply that CoA is not the preferred substrate. The activity of NudC with three structurally distinct coenzymes demonstrated here reinforces the lack of specificity previously seen by Frick et al. (1995), and provides another example of the substrate ambiguity attributed to Nudix hydrolases [27].

The demonstration of two enzymes with substrate overlap is not new to *E. coli*, especially not in the Nudix superfamily [2], thus the existence of two enzymes that coevolved with a shared CoA-hydrolyzing activity in *E. coli* is not farfetched. While NudL maintains the specificity for CoA- and CoA-related molecules with its CoA motif (Fig. S7), NudC likely plays the role of a general coenzyme hydrolase. The slower kinetic constants determined for these enzymes compared to eukaryote orthologs are also plausible given the lack of intracellular compartmentalization in bacteria. Indeed, eukaryotic CoA hydrolases have much higher turnover numbers [7, 8, 11], but these proteins are localized within the peroxisome, and are thereby separated from the cellular stores of CoA in the mitochondria. The existence of such a highly active enzyme within a bacterial cell could be detrimental to its survival considering the importance of CoA to so many cellular functions. Given their relative transcript abundance [2], these enzymes, therefore, are likely kept under tight regulation at the translational level to avoid interference with normal metabolic activity. Furthermore, the low solubility of NudL may be a form of regulation itself, where *in vivo* activity depends on cooperative interactions with other proteins such as chaperones.

Recent studies have determined that NudC prefers NAD-RNA over NADH [20] with a purported primary role of NudC of NAD-RNA decapping. We’ve demonstrated here along with others [19] that NudC is also capable of cleaving CoA-RNA, but this activity is a kinetic afterthought in comparison to NAD and NAD-RNA. The function and capping mechanisms for NAD- and CoA-RNA are currently unknown, but the two caps likely maintain different functions, and therefore their degradation could be catalyzed by different enzymes. The activity of murine Nudt7 with CoA-thioesters [11] and CoA-RNA (Fig. 4) make its ortholog NudL a likely candidate for a specific CoA-RNA decapping enzyme in *E. coli* (Fig. S7). Despite sequence similarities with Nudt7, CoA-RNA decapping with MBP-NudL could not be observed under our experimental conditions. It’s unclear if such activity was impeded by the presence of MBP, but the fusion was still active against oxidized CoA indicating it could accommodate molecules extending from the phosphopantetheine end of the original CoA structure such as CoA thioesters. The possibility also exists that the rate of CoA-RNA degradation by NudL was too slow to be observed on the same scale as the much faster enzymes NudC and Nudt7. Considering that the RNA extends from the 3’-OH of the adenosine in CoA, it is possible that CoA-RNA presents a steric hindrance to efficient phosphohydrolysis by the bulky MBP-NudL fusion not observed with CoA-dimers extending from the opposite end of the molecule.

The slow kinetics and poor *in vitro* solubility imply that NudL may have little biological significance aside from its specificity for CoA and CoA-derivatives, especially in comparison to the more efficient NudC. An enzyme capable of specific coenzyme degradation, however, would be advantageous considering the bacterial intracellular environment lacks compartmentalization. A highly active NudC within *E. coli* would result in the hydrolysis of NAD^+^, NADH, FAD, CoA, NAD-RNA, and CoA-RNA with substrate affinity acting as the only form of discrimination. The unregulated degradation of coenzymes would result in a depressed energetic state and ultimately lead to cell death if unchecked. In contrast, the cell could employ NudL to specifically cleave CoA-related structures at a slower rate without affecting cellular stores of NAD or FAD, thereby providing a more controlled degradation pathway. While there is no evidence that a lack of CoA degradation is lethal, CoA synthesis is essential to survival [28], and Nudix-catalyzed CoA phosphohydrolysis is a quicker route to generate the precursor pPan than either the CoA pathway or β-oxidation. The yeast ortholog of NudL was hypothesized to function in the clearance of oxidized, inactive CoA [8], and considering NudL is similarly active against oxidized CoA, one would expect a NudL knockout to have slower growth due to such metabolic strain.

## 5. Conclusions

Here we present evidence of two separate CoA-degrading enzymes that, to our knowledge, are the first characterized enzymes to carry out this activity in *E. coli*. Both are members of the Nudix superfamily of phosphohydrolases and carry out similar phosphohydrolysis mechanisms with different specificity and kinetics. The variety of substrates hydrolyzed by NudC implies its intracellular role as a high activity general coenzyme hydrolase. NudL represents the opposite side of the coin with specificity for CoA-related molecules and low activity. These findings open the door for further studies on the roles of these enzymes as a cooperative CoA- and CoA-RNA regulation system. Furthermore, investigations into the solubility of the native NudL may reveal kinetic constants more comparable to its orthologs as well as the proposed capability for CoA-RNA decapping, providing more insight to a multi-faceted system for CoA and CoA-RNA regulation.

## Acknowledgements

We would like to acknowledge the Mississippi INBRE program supported by the National Institutes of Health for using its facilities.

## Reference

[1] A.S. Mildvan, Z. Xia, H.F. Azurmendi, V. Saraswat, P.M. Legler, M.A. Massiah, S.B. Gabelli, M.A. Bianchet, L.W. Kang, L.M. Amzel, Structures and mechanisms of Nudix hydrolases, Arch Biochem Biophys, 433 (2005) 129–143

[2] A.G. McLennan, The Nudix hydrolase superfamily, Cell Mol Life Sci, 63 (2006) 123–143.

[3] J.R. Srouji, A. Xu, A. Park, J.F. Kirsch, S.E. Brenner, The evolution of function within the Nudix homology clan, Proteins, 85 (2017) 775–811.

[4] M.J. Bessman, D.N. Fricks, S.F. O’Handley, The MutT Proteins or “Nudix” Hydrolases, a Family of Versatile, Widely Distributed, “Housecleaning” Enzymes, J Biol Chem, 271 (1996) 25059–25062.

[5] D.N. Frick, M.J. Bessman, Cloning, Purification, and Properties of a Novel NADH Pyrophosphatase, J Biol Chem, 270 (1995) 1529–1534.

[6] W. Xu, C.A. Dunn, M.J. Bessman, Cloning and characterization of the NADH pyrophosphatases from Caenorhabditis elegans and Saccharomyces cerevisiae, members of a Nudix hydrolase subfamily, Biochem Bioph Res Co, 273 (2000) 753–758.

[7] S.R. AbdelRaheim, A.G. McLennan, The Caenorhabditis elegans Y87G2A.14 Nudix Hydrolase is a Peroxisomal Coenzyme A Diphosphatase, BMC Biochem, 3 (2002) 5.

[8] J.L. Cartwright, L. Gasmi, D.G. Spiller, A.G. McLennan, The Saccharomyces cerevisiae PCD1 gene encodes a peroxisomal nudix hydrolase active toward coenzyme A and its derivatives, J Biol Chem, 275 (2000) 32925–32930.

[9] L.W. Kang, S.B. Gabelli, M.A. Bianchet, W.L. Xu, M.J. Bessman, L.M. Amzel, Structure of a Coenzyme A Pyrophosphatase from Deinococcus radiodurans: a Member of the Nudix Family, J Bacteriol, 185 (2003) 4110–4118.

[10] T. Kupke, J.A. Caparrós-Martín, K.J. Malquichagua Salazar, F.A. Culiáñez-Macià, Biochemical and physiological characterization of Arabidopsis thaliana AtCoAse: a Nudix CoA hydrolyzing protein that improves plant development, Physiol Plantarum, 135 (2009) 365–378.

[11] L. Gasmi, G. McLennan, The Mouse Nudt7 Gene Encodes a Peroxisomal Nudix Hydrolase Specific for Coenzyme A and its Derivatives, J Biochem, 357 (2001) 33–38.

[12] S. Jackowski, C.O. Rock, Regulation of Coenzyme A Biosynthesis, J Bacteriol, 148 (1981) 926–932.

[13] S. Jackowski, C.O. Rock, Consequences of Reduced Intracellular Coenzyme A Content in Escherichia coli, J Bacteriol, 166 (1986) 866–871.

[14] S. Jackowski, C.O. Rock, Metabolism of 4’-Phosphopantetheine in Escherichia coli, J Bacteriol, 158 (1984) 115–120.

[15] R. Leonardi, Y.M. Zhang, C.O. Rock, S. Jackowski, Coenzyme A: back in action, Prog Lipid Res, 44 (2005) 125–153.

[16] Y.G. Chen, W.E. Kowtoniuk, I. Agarwal, Y. Shen, D.R. Liu, LC/MS analysis of cellular RNA reveals NAD-linked RNA, Nat Chem Biol, 5 (2009) 879–881.

[17] W.E. Kowtoniuk, Y. Shen, J.M. Heemstra, I. Agarwal, D.R. Liu, A chemical screen for biological small molecule-RNA conjugates reveals CoA-linked RNA, P Natl Acad Sci USA, 106 (2009) 7768–7773.

[18] H. Cahova, M.L. Winz, K. Hofer, G. Nubel, A. Jaschke, NAD captureSeq indicates NAD as a bacterial cap for a subset of regulatory RNAs, Nature, 519 (2015) 374–377.

[19] J.G. Bird, Y. Zhang, Y. Tian, N. Panova, I. Barvik, L. Greene, M. Liu, B. Buckley, L. Krasny, J.K. Lee, C.D. Kaplan, R.H. Ebright, B.E. Nickels, The mechanism of RNA 5’ capping with NAD+, NADH and desphospho-CoA, Nature, (2016).

[20] K. Hofer, S. Li, F. Abele, J. Schlotthauer, J. Grawenhoff, J. Du, D.J. Patel, A. Jaschke, Structure and function of the bacterial decapping enzyme NudC, Nat Chem Biol, (2016).

[21] F. Huang, Efficient incorporation of CoA, NAD and FAD into RNA by in vitro transcription, Nucleic Acids Res, 31 (2003) e8.

[22] T.M. Coleman, F. Huang, RNA-Catalyzed Thioester Synthesis, Chem Biol, 9 (2002) 1227–1236.

[23] M.P. Malakhov, M.R. Mattern, O.A. Malakhova, M. Drinker, S.D. Weeks, T.R. Butt, SUMO fusions and SUMO-specific protease for efficient expression and purification of proteins, J Struct Func Genom, 5 (2004) 75–85.

[24] G.L. Rosano, E.A. Ceccarelli, Recombinant protein expression in Escherichia coli: advances and challenges, Front Microbiol, 5 (2014) 172.

[25] D. Esposito, D.K. Chatterjee, Enhancement of soluble protein expression through the use of fusion tags, Curr Opin Biotech, 17 (2006) 353–358.

[26] A. Ray, B.A. Beaupre, G.R. Moran, D.N. Frick, The E. coli and Human Nudix hydrolases NudC and NUDT12 cleave damaged NADH, FASEB J, 31 (2017) 606.11.

[27] A.G. McLennan, Substrate ambiguity among the nudix hydrolases: biologically significant, evolutionary remnant, or both?, Cell Mol Life Sci, 70 (2013) 373–385.

[28] C.J. Balibar, M.F. Hollis-Symynkywicz, J. Tao, Pantethine rescues phosphopantothenoylcysteine synthetase and phosphopantothenoylcysteine decarboxylase deficiency in Escherichia coli but not in Pseudomonas aeruginosa, J Bacteriol, 193 (2011) 3304–3312.

