## Supplemental Information for "The *E. coli* NudL enzyme is a Nudix hydrolase that cleaves CoA and its derivatives"

**Supplementary Material**

Figure S1. **Purification of MBP-NudL**. SDS-PAGE of MBP-NudL purification showing the soluble cell lysate (S), column flow-through (F), final wash fraction (W) and 1 mL elution fractions using 20 mM maltose (E1 – E5).

Figure S2. **Solubility of SUMO-NudC**. SDS-PAGE of SUMO-NudC comparing fractions of uninduced cell lysate (U), induced cell lysate (I), soluble cell lysate (S) and pelleted cell lysate (P).

Figure S3. **Purification of SUMO-NudC**. SDS-PAGE of SUMO-NudC purification showing first 1 mL wash fraction with 20 mM imidazole (W1), final 1 mL wash fraction with 20 mM imidazole (W8), and 1 mL elution fractions with 300 mM imidazole (E1 – E7).

Figure S4. **Thrombin Cleavage of MBP-NudL**. SDS-PAGE of thrombin cleavage for removal of MBP fusion from MBP-NudL.

Figure S5. **$NAD^+$ cleavage by SUMO-NudC**. HPLC chromatogram showing separation of NMN and AMP from $NAD^+$ during NudC hydrolysis. All three species have absorbance at 260 nm. NMN and AMP coelute as a single peak since both molecules contain a single phosphate that is distinct from two phosphate-containing $NAD^+$.

Figure S6. **FAD cleavage by SUMO-NudC**. **A)** Chemistry of FAD hydrolysis by NudC. **B)** HPLC chromatogram showing separation of FMN and AMP from FAD during NudC hydrolysis.

Figure S7. **Amino acid sequence comparison of known CoA hydrolyzing Nudix enzymes.** Comparison of 8 proven CoA-hydrolyzing Nudix enzymes and 1 hypothesized based on sequence similarity (Human NUDT7) to the consensus sequence denoting a Nudix box downstream of a CoA motif. Bold underlined amino acids highlight those identical to the consensus sequence. All examples except for NudC contain a CoA motif, indicating their specificity for CoA hydrolysis. Figure adapted and modified from Gasmi & McLennan (2001) to show larger diversity across different phyla.

Figure S1. **Purification of MBP-NudL**.

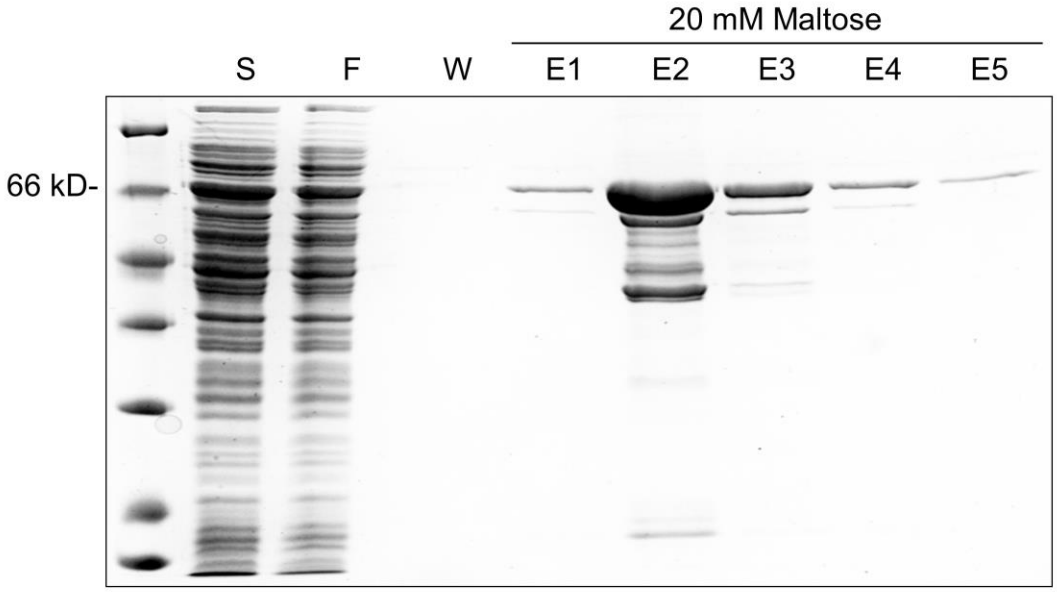

Figure S2. **Solubility of SUMO-NudC**.

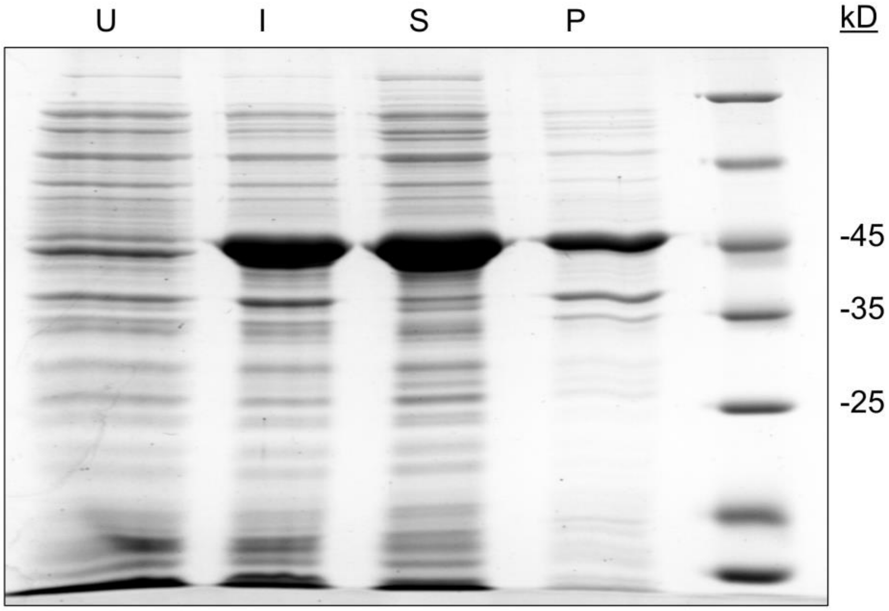

Figure S3. **Purification of SUMO-NudC**.

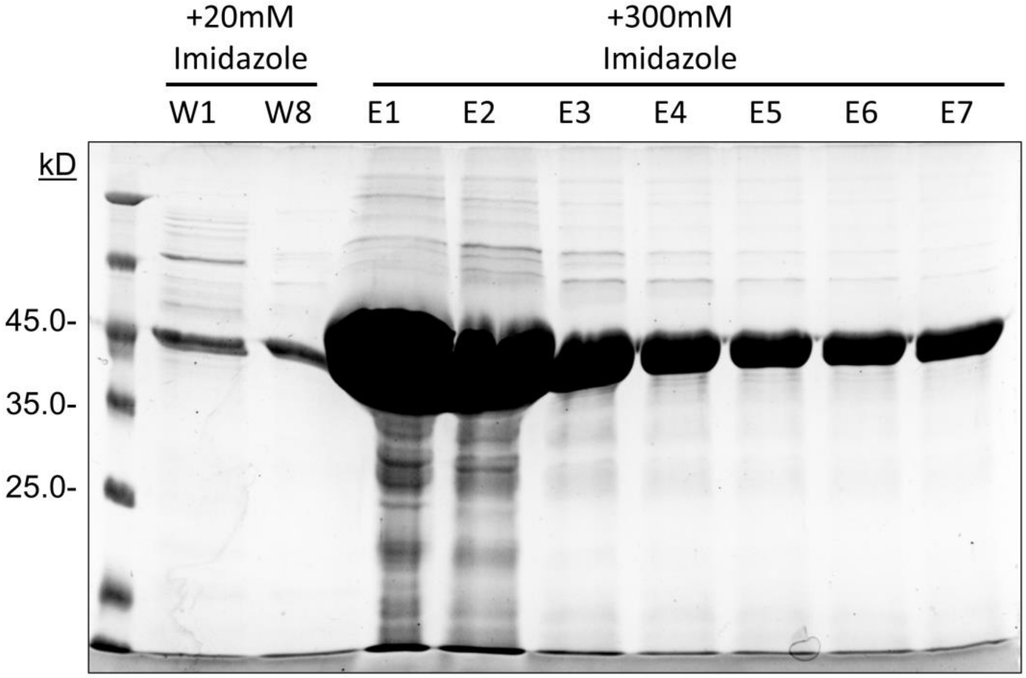

Figure S4. **Thrombin Cleavage of MBP-NudL**.

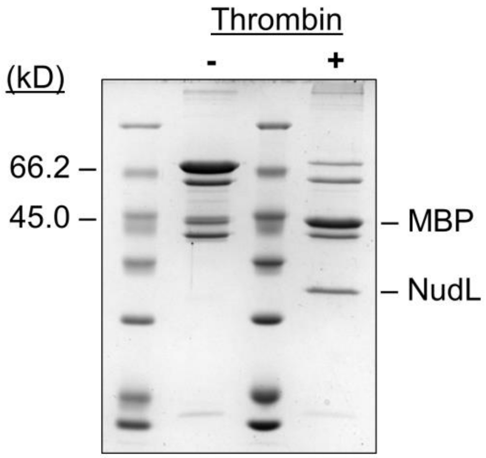

Figure S5. **NAD$^+$ cleavage by SUMO-NudC**.

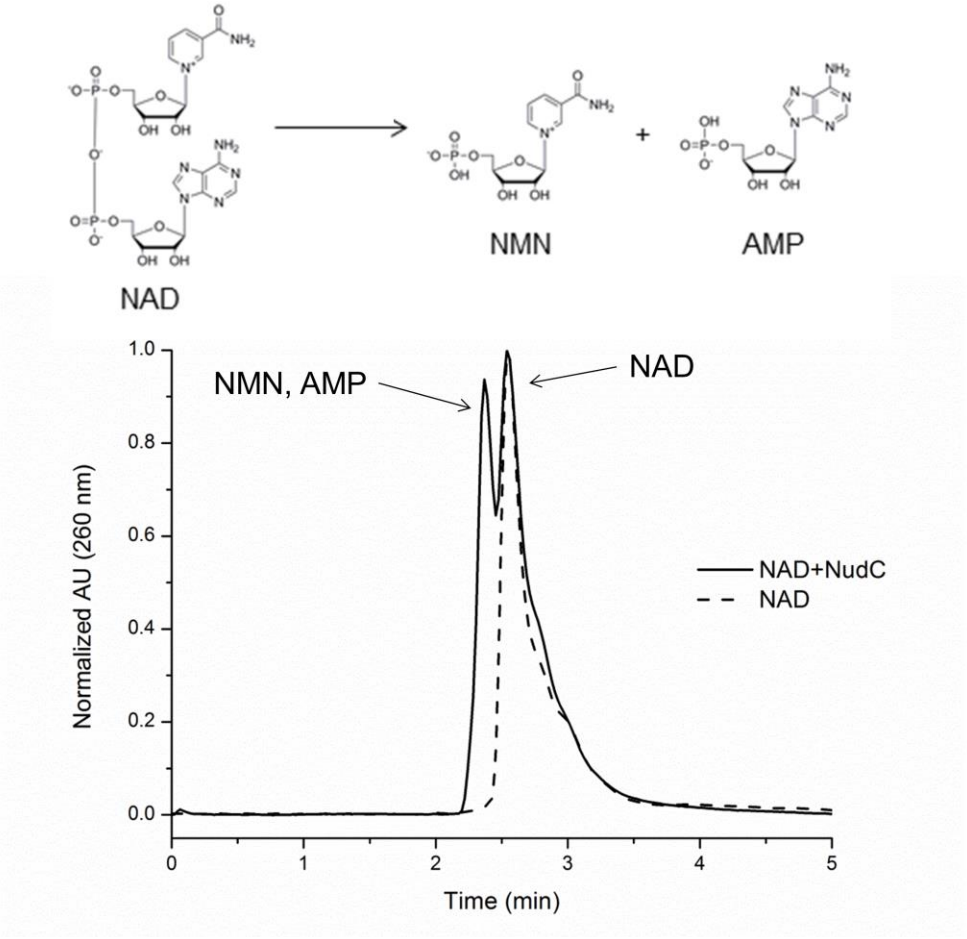

Figure S6. **FAD cleavage by SUMO-NudC**.

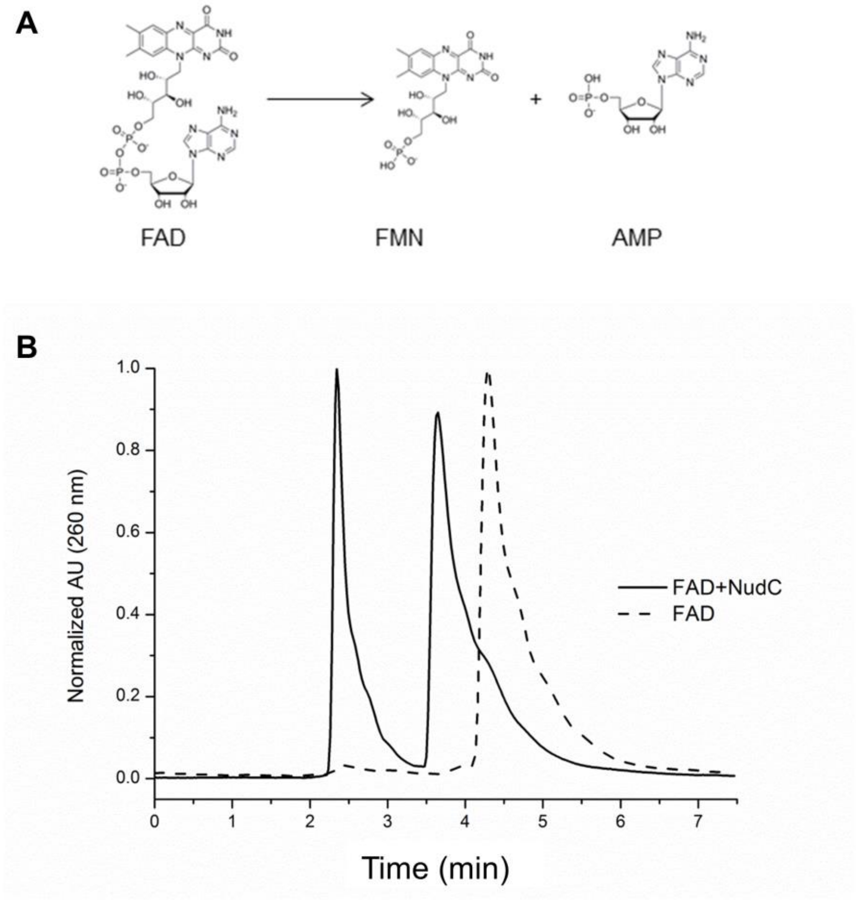

Figure S7. **Amino acid sequence comparison of known CoA hydrolyzing Nudix enzymes.**

```
                         CoA Motif (UPF0035)
                                             Nudix Box (PS00893)
Consensus                xLLTxRSxxxR-xxxGxxxFPGGxxxxxE--xxxxxAxRExxEExGUxxx
Mouse Nudt7α             LMFTVRSDKLK-REPGEVCFPGGKRDPVDT-DDTATALREAQEEVGLHPH
E.coli NudL              LLLTQRSIHLR-KHAGQVAFPGGAVDDTDA-SAIAAALREAEEEVAIPPS
E.coli NudC              RDDSILLAQHTRHRNGVHTVLAGFVEVGET-LEQAVA-REVMEESGIKVK
C.elegans NDX8           VLLTKRSIHLR-SHRGEVCFPGGRMDPDGE-TTTETALRETFEEIGVNAE
S.cerevisiae Pcd1p       VLLTKRSRTLR-SFSGDVSFPGGKADYFQE-TEESVARREE-EEIGLPHD
Human NUDT7              LLFTVRSEKLR-RAPGEVCFPGGKRDPTDM-DDAATALREAQEEVGLRPH
D.radiodurans 1184       VLLTVRSSELP-THKGQISFPGGSLDAGET--PTQAALREAQEEVALDPA
A.thaliana NUDX11        VILTKRSTTLS-SHPGEVALPGGKRDQEDK-DDIATALREAREEIGLDPS
A.thaliana NUDX15        VILTKRSSKLS-THSGEVSLPGGKAEEDDK-DDGMTATREAEEEIGLDPS
```
